# Mechanism of formin-mediated filament nucleation from profilin-actin

**DOI:** 10.1101/2025.05.09.653129

**Authors:** Mark E. Zweifel, Naomi Courtemanche

## Abstract

Formins direct the assembly of unbranched actin filaments through dynamic interactions with the abundant, actin-binding protein profilin. Although the process of formin-mediated filament elongation is well characterized, the mechanism by which formins nucleate actin filaments is poorly understood. In this study, we used *in vitro* reconstitution assays and kinetic modeling to dissect filament nucleation mediated by the *S. cerevisiae* formin Bni1p. We found that formins assemble filament nuclei by sequentially binding three actin monomers in a process that takes place in parallel to spontaneous actin assembly. Whereas the formin FH2 domain preferentially binds actin monomers, the flexible FH1 domain facilitates *de novo* filament formation by binding and transferring profilin-actin complexes to a free FH2 dimer. FH1-mediated delivery stimulates filament nucleation in a distance- and sequence-dependent manner. The architectures of FH1 domains therefore maximize the efficiency of formin-mediated polymerization through optimized engagement of each of their profilin-binding sites.

## Introduction

Actin polymerization drives the assembly of dynamic cytoskeletal networks that facilitate cell growth, motility and division (1-3). Polymerization proceeds via an initial nucleation step in which a minimally-stable, trimeric actin “nucleus” is assembled (4-6). The addition of a fourth monomer converts the nucleus into a filament that elongates through continued actin binding at both ends (4, 7). As the rate-limiting step in polymerization, nucleation is tightly regulated by a diverse set of actin-binding proteins (8, 9).

Formins are a family of actin assembly factors that stimulate the nucleation and barbed-end elongation of unbranched actin filaments (10, 11). Filaments assembled by formins are incorporated into higher-order structures, including cytokinetic rings, filopodia, actin cables, and stress fibers (11-13). Formins bind actin via their dimeric formin homology 2 (FH2) domains, which encircle filament nuclei and barbed ends (14-16). FH2 dimers are proposed to promote the assembly of filament nuclei by binding two actin monomers before facilitating the addition of a third monomer (17). Following nucleation, each FH2 subunit within the dimer takes alternating steps onto incoming actin monomers, thereby retaining the formin at the barbed end of the elongating filament (18).

In cells, formins overcome energetic barriers to polymerization through dynamic interactions with the abundant, actin-binding protein profilin (19, 20). Profilin-actin complex formation inhibits spontaneous filament nucleation and pointed-end elongation but permits filament growth through repeated cycles of profilin-actin addition and profilin dissociation at the barbed end (21-23). Once polymerized, filamentous actin subunits undergo conformational flattening, which weakens their affinity for profilin by at least two orders of magnitude (24, 25). This change in affinity releases profilin from the barbed end, thereby enabling subsequent actin binding (25).

In addition to its interactions with actin, profilin binds polyproline sequences encoded in the structurally flexible formin FH1 domain (26). These interactions enable diffusion-limited, FH1-mediated delivery of profilin-actin complexes to FH2-bound filament barbed ends (18, 27). This delivery step speeds filament elongation beyond the rate achieved through direct, FH1-independent binding of actin and profilin-actin to the barbed end (28, 29). The magnitude of this effect on elongation depends on the number of polyproline tracts contained by the FH1 domain, as well as each tract’s location and affinity for profilin (27, 30).

Pioneering studies of filament assembly mediated by the *S. cerevisiae* formin Bni1p revealed that FH2-directed nucleation slows in the presence of profilin (31-33). Whereas the FH1 domain plays no role in nucleation in the absence of profilin, a construct encoding the FH1 and FH2 domains of Bni1p generates faster polymerization in the presence of profilin than does the FH2 domain on its own (33). This finding is consistent with a central role for the FH1 domain in formin-mediated nucleation. However, although the interactions that occur between formins and profilin during filament elongation have been extensively characterized (18, 30, 34), the mechanism by which formins promote actin filament nucleation in the presence of profilin remains poorly understood. As such, it is not known how formins generate filament nuclei using profilin-actin complexes, and how this activity influences the overall process of polymerization.

In this study, we systematically dissected the mechanism of filament nucleation mediated by Bni1p. Using a deterministic kinetic model to fit experimental polymerization time courses, we found that FH2 dimers can bind individual monomers sequentially and do not require pre-assembled actin dimers during nucleation. Whereas the FH2 domain preferentially binds actin monomers, the FH1 domain facilitates the assembly of filament nuclei by delivering profilin-actin complexes to a free FH2 dimer. The overall rate of formin-mediated polymerization increases as the distance separating the FH2 domain and the location at which profilin binds the FH1 domain increases, revealing an inverse-dependence on the rate of FH1-mediated delivery. This effect arises from differential, position-specific modulation of the rates of filament nucleation and elongation, suggesting that the FH1 domain contributes differently to each process. Sequence variations confer specialized nucleation and elongation activities to each profilin-binding polyproline tract within an FH1 domain. Collectively, the sequence architectures of FH1 domains maximize the efficiency of formin-mediated polymerization through optimized engagement of each of their profilin-binding sites during polymerization.

## Results

To investigate the mechanism of formin-mediated actin filament nucleation, we analyzed the polymerization activity of the *S. cerevisiae* formin Bni1p. This formin has robust nucleation and elongation properties (19, 31, 35), which provide a large dynamic range of activity that facilitates detailed characterization. To quantify Bni1p-mediated nucleation, we measured the fluorescence emission of 2 µM actin (20% pyrene-labeled) over the course of bulk polymerization reactions performed in the presence of a range of concentrations of the formin’s FH2 domain (called “Bni1p FH2”). The fluorescence signal in these assays provides a direct measurement of the concentration of polymerized actin (36, 37). As previously reported, Bni1p FH2 increases the polymerization rate—which is measured via the application of linear fits to the time courses at the point at which half the actin is polymerized—in a concentration-dependent manner (Figure 1A) (32, 33, 38). Filaments bound by Bni1p FH2 elongate approximately half as fast as filaments with free barbed ends (19), so this increase in the polymerization rate is consistent with an increase in filament nucleation.

**Figure 1.**
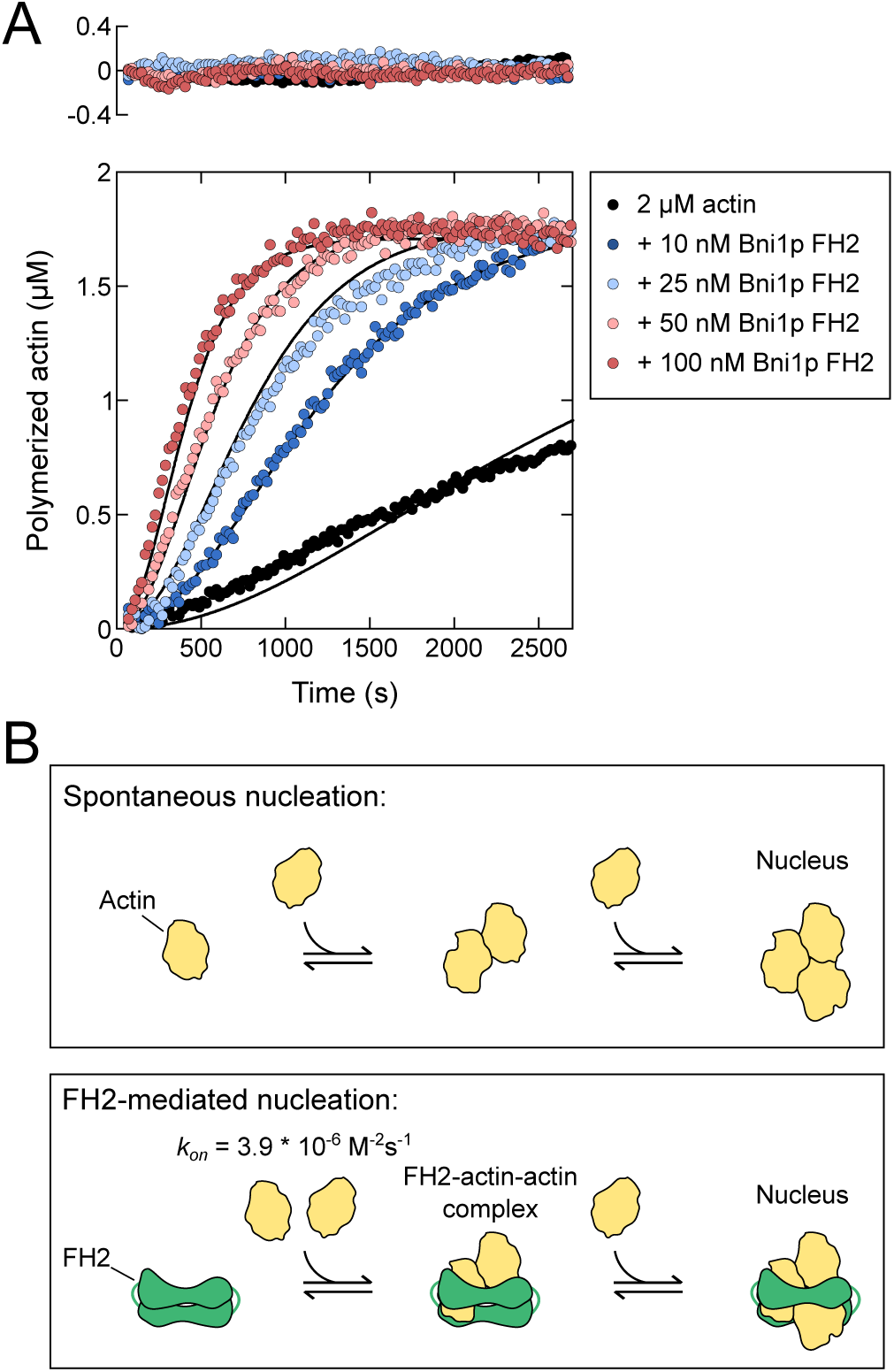
The FH2 domain stimulates nucleation by creating stable actin dimers. (A) Representative time courses of spontaneous polymerization of 2 µM actin (20% pyrene-labeled) in the presence of a range of concentrations of Bni1p FH2. For clarity, every second data point is plotted. Solid lines are fits to the data using our kinetic model for FH2-mediated nucleation. Upper panel shows residuals for each fit. (B) Schematic depicting spontaneous and FH2-mediated actin filament nucleation. Actin monomers and an FH2 dimer are depicted in yellow and green.

To extract kinetic parameters for nucleation, we used a deterministic computational model to apply global fits to our polymerization time courses. Our model accounts for interactions among actin monomers, actin nuclei, filament ends, and FH2 dimers, and enables independent quantification of filament nucleation and elongation over time (Supplemental Table S1) (35). To account for the stability of FH2 dimers, which do not undergo assembly or disassembly over the period of our polymerization reactions (14), our model considers each FH2 dimer to function as a single molecular species. Consistent with previous studies, our model considers spontaneous (i.e., formin-independent) nucleation to occur via an initial actin dimerization step, which is followed by the addition of a third actin monomer to establish a filament “nucleus” (Figure 1B) (5, 37). Addition of a fourth monomer converts the nucleus into a filament, which elongates at both ends at distinct and well-characterized rates (7). FH2 dimers promote nucleation by binding two actin monomers and facilitating the addition of a third monomer to establish a formin-bound nucleus (Figure 1B) (20, 33, 35).

The global fits of our experimental data produce randomly-distributed residuals (Figure 1A, upper panel), indicating that spontaneous and FH2-mediated actin assembly are well described by our model. Our fits yield association rates governing the interactions of FH2 dimers with the three actin monomers composing filament nuclei. The rate at which FH2 dimers bind the first actin monomer is faster than the rate at which the second monomer is bound, owing to reductions in the number of available actin-binding sites and the rate of FH2 diffusion following the first actin-binding event. As a result, a simplified model in which an FH2 dimer binds two actin monomers simultaneously before binding a third monomer also generates good fits to our data (20, 33). This model enables the quantification of a trimeric association rate describing the assembly of the FH2-actin-actin species without requiring any assumptions regarding the rates at which each independent binding event occurs. As this simplified model produces the best-defined set of kinetic parameters, we use it in this study to describe the initial binding step that occurs during FH2-mediated nucleation.

To assess the robustness of our model, we used two widely divergent sets of published rates of spontaneous actin nucleation as starting points for our fitting (Supplemental Table S1) (5, 37). Both sets of rates produce identical parameters defining the interactions between FH2 dimers and actin monomers during nucleus formation (Figure 1B). These results suggest that FH2-mediated nucleation does not require actin self-oligomerization.

### The formin FH1 domain enhances nucleation of profilin-actin

Binding of profilin to actin monomers inhibits spontaneous filament nucleation and slows FH2-mediated polymerization (21-23, 33). To quantify these impacts on polymerization, we collected time courses of bulk actin assembly in the presence of a range of concentrations of *S. cerevisiae* profilin. We compared rates of polymerization mediated by Bni1p FH2 to those generated by a construct encoding both the FH1 and FH2 domains (called “Bni1p FH1FH2”). As previously reported, the addition of 5 µM profilin results in near-complete inhibition of actin assembly in the absence of formin (Figure 2A, black data) (21, 23). At this concentration of profilin, Bni1p FH2 stimulates polymerization at approximately 40% of the rate that is generated by this formin in the absence of profilin (Figure 2A and 2B, green data). FH2-mediated polymerization slows further as the concentration of profilin increases. In the presence of 25 µM profilin, nearly all monomers are profilin-bound (39), and the polymerization rate is 85% slower than that observed in reactions lacking profilin. Binding of profilin slows the elongation of FH2-bound and free filaments by ∼50% at 25 µM profilin (25, 29), thereby increasing the critical concentration and reducing the equilibrium concentration of filamentous actin (23). The 85% reduction in the bulk polymerization rate at this concentration of profilin indicates that nucleation also slows in the presence of profilin. Collectively, these data suggest that the FH2 domain can bind and utilize profilin-actin to assemble filament nuclei, but this process is relatively inefficient.

**Figure 2.**
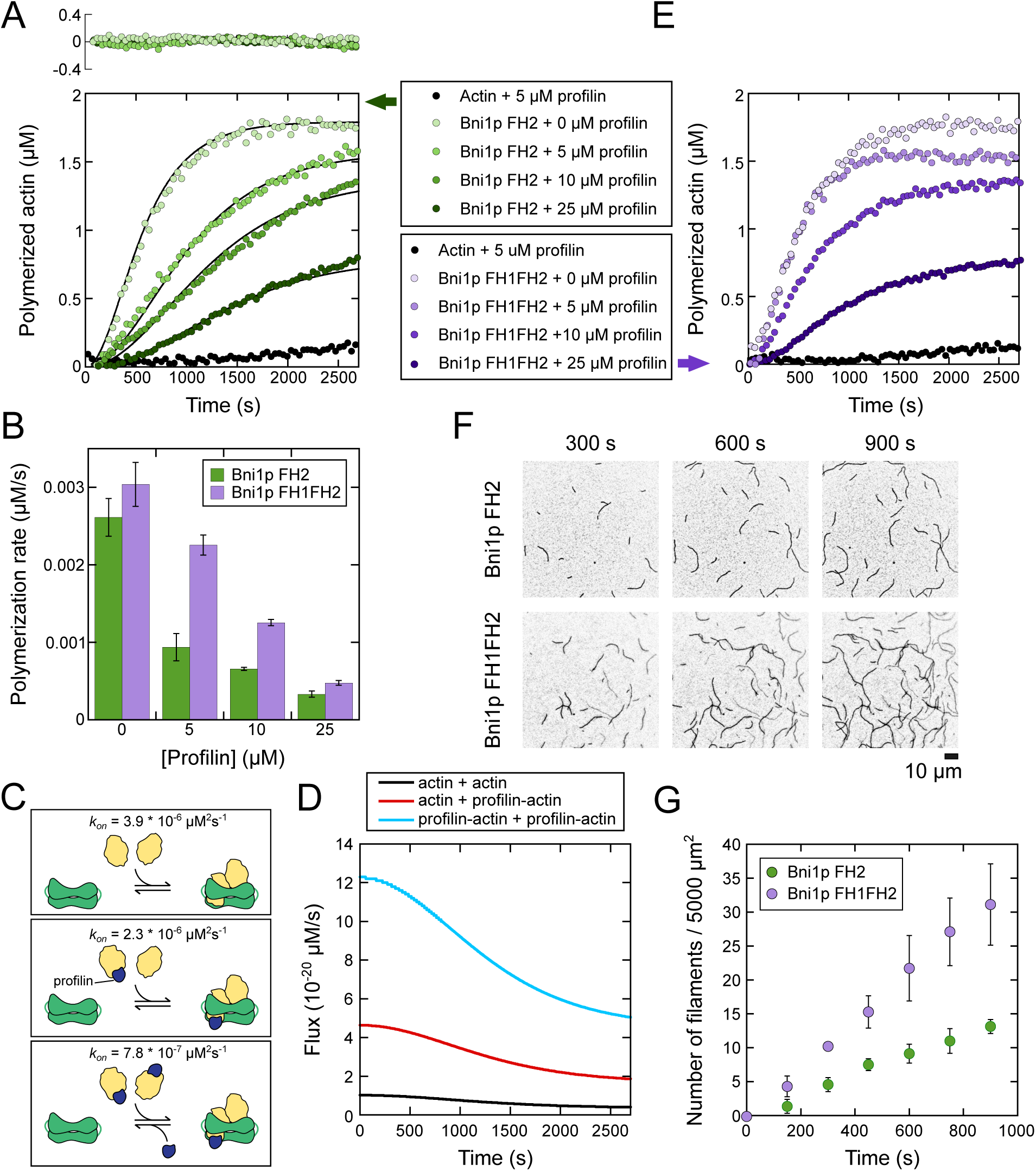
The formin FH1 domain enhances nucleation of profilin-actin. (A) Representative time courses of spontaneous polymerization of 2 µM actin (20% pyrene-labeled) in the presence of 50 nM Bni1p FH2 and a range of concentrations of *Sc* profilin. The dataset collected in the absence of profilin is identical to that shown in Figure 1A. For clarity, every fourth data point is plotted. Solid lines are fits to the data using our kinetic model for FH2-mediated nucleation in the presence of profilin. Upper panel shows residuals for each fit. (B) Polymerization rates obtained from the slope of the change in fluorescence signal where half of the actin is polymerized in reactions containing 50 nM Bni1p FH2 (green) or Bni1p FH1FH2 (purple) and a range of concentrations of profilin. Error bars are standard deviations of the mean polymerization rate calculated from 3 independent assays. (C) Schematic depicting three pathways to assemble the FH2-actin-actin complex (green and yellow) in the presence of profilin (dark blue). Association rates extracted from kinetic fits to our experimental data are indicated for each pathway. (D) Flux through three parallel pathways of FH2-actin-actin complex assembly in the presence of profilin plotted as a function of time. Concentrations of FH2-actin-actin complexes per second were obtained from global fits to experimental time courses as depicted in panel A. The FH2-actin-actin complex can be formed via FH2 binding of two actin monomers (black line), one actin monomer and one profilin-actin complex (red line), or two profilin-actin complexes (blue line). (E) Representative time courses of spontaneous polymerization of 2 µM actin (20% pyrene-labeled) in the presence of 50 nM Bni1p FH1FH2 and a range of concentrations of *Sc* profilin. For clarity, every fourth data point is plotted. (F) Time series of inverted fluorescence micrographs of filaments assembled by 5 nM Bni1p FH2 or Bni1p FH1FH2 in the presence of 0.75 µM actin monomers (33% Oregon Green-labeled) and 1 µM *Sc* profilin. Micrographs were collected by TIRF microscopy. The scale bar is 10 µm. (G) Number of filaments visualized per 5,000 µm^2^ in polymerization assays containing 0.75 µM actin monomers (33% Oregon Green-labeled), 1 µM *Sc* profilin, and 5 nM Bni1p FH2 or Bni1p FH1FH2 as shown in panel E. Error bars are standard deviations of the mean number of filaments visualized in 3 independent assays.

We applied global fits to our polymerization time courses using our kinetic model to quantify the efficiency of FH2-mediated nucleation in the presence of profilin (Figure 2A and C). Steric constraints dictate that profilin can bind only the terminal actin subunit at the barbed end of a filament (25). Therefore, we assumed that profilin must dissociate from any profilin-actin complex that is bound by the FH2 domain before the next actin subunit can bind. Our fitting revealed that FH2 dimers bind two actin monomers nearly twice as fast as they bind one actin monomer and one profilin-actin complex, and five times faster than they bind two profilin-actin complexes. In the presence of 25 µM profilin, 94% of formin-bound filaments are nucleated via an initial binding event involving at least one profilin-actin complex (Figure 2D).

Whereas Bni1p FH2 and Bni1p FH1FH2 possess identical polymerization activities in the absence of profilin, Bni1p FH1FH2 mediates faster actin assembly at all concentrations of profilin (Figure 2B and E). To determine whether this enhanced polymerization activity stems solely from the faster filament elongation rate produced by Bni1p FH1FH2, we visualized polymerization mediated by both constructs in the presence of 1 µM profilin using total internal reflection fluorescence (TIRF) microscopy (Figure 2F). At this concentration of profilin, filaments bound by Bni1p FH1FH2 elongate approximately four times faster than filaments bound by Bni1p FH2 (19, 20, 29). Micrographs collected at regular intervals over the course of polymerization reveal the assembly of approximately twice as many filaments in reactions containing Bni1p FH1FH2 than in reactions containing an identical concentration of Bni1p FH2 (Figure 2G). Therefore, the FH1 domain stimulates actin polymerization in the presence of profilin by enhancing both nucleation and elongation.

### Simplifying the interactions of FH1 domains with profilin

The increase in the number of filaments assembled by Bni1p when its FH1 domain is present suggests that FH1-mediated delivery facilitates the binding of FH2 dimers to profilin-actin complexes during nucleation. FH1 domains encode multiple polyproline tracts, which bind profilin with micromolar-to-millimolar affinity, depending on their length (26, 40). Each of Bni1p’s four polyproline tracts binds and delivers profilin-actin complexes to the barbed end during filament elongation (30). Therefore, all four tracts may also contribute to nucleation.

To dissect the role of the FH1 domain in Bni1p-mediated nucleation, we sought to simplify the formin’s interactions with profilin. We did so by combining the sequential reactions in which profilin first binds an actin monomer and then binds the FH1 domain into a single binding event (Figure 3A). We purified a series of variants of Bni1p FH1FH2 in which profilin is covalently tethered to the FH1 domain. In each variant, we replaced a single polyproline tract with the amino acid sequence of *S. cerevisiae* profilin (Figure 3B) (34). Because profilin’s N-and C-termini are both located on its polyproline-binding surface, this covalent linkage positions profilin in a manner that mimics the orientation it adopts when bound to a polyproline tract (41). We substituted the three remaining polyproline tracts with poly-Gly/Ser sequences to retain the structural flexibility of the FH1 domain. Each chimeric formin-profilin construct therefore facilitates direct binding of an actin monomer to a single site in the FH1 domain with uniform affinity. We named our variants “PRF_xx_FH2”, where “xx” refers to the number of amino acids separating profilin from the FH2 domain (34).

**Figure 3.**
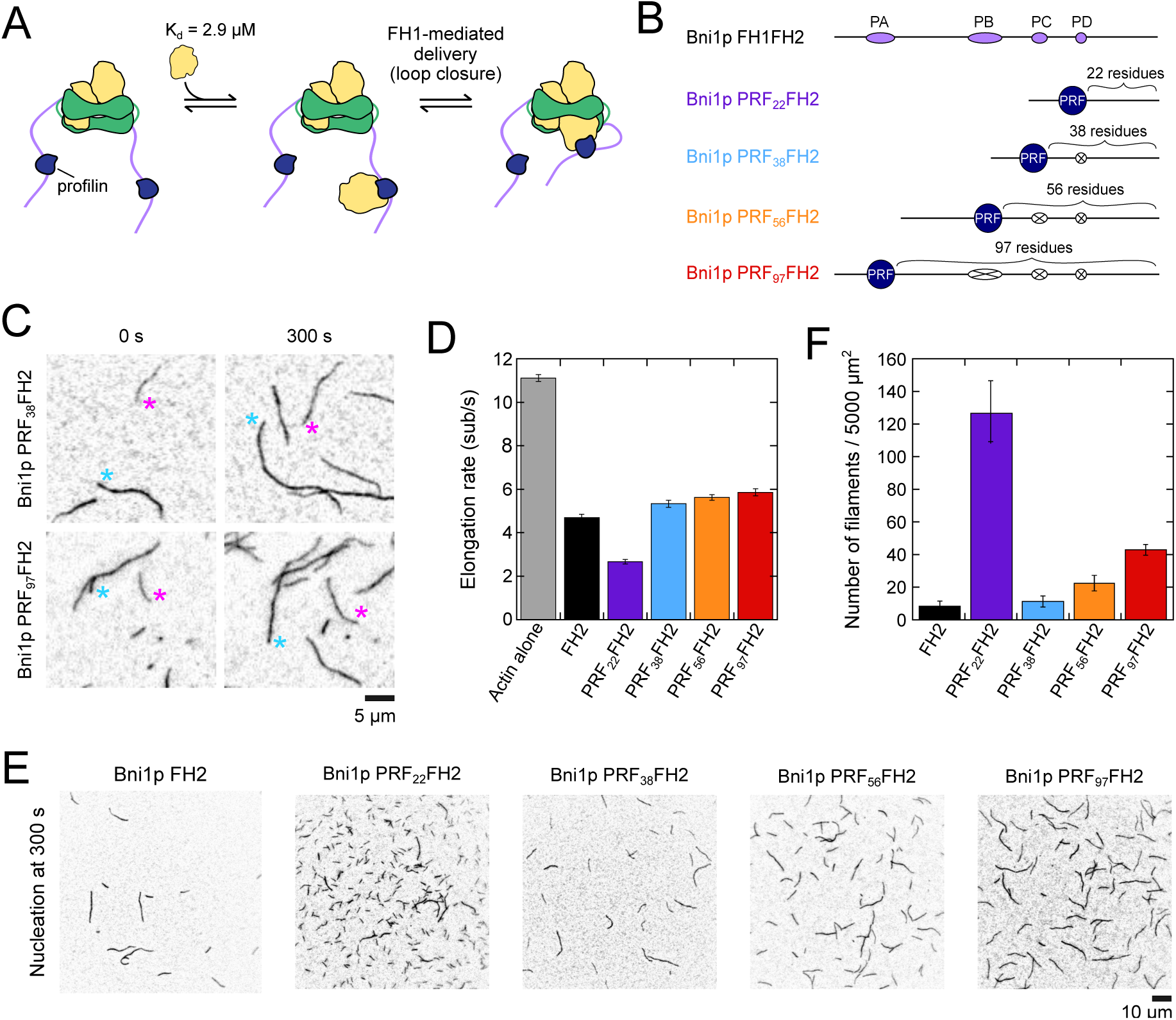
Simplifying the interactions of FH1 domains with profilin. (A) Schematic depicting the binding and delivery of an actin monomer (yellow) by a formin whose FH1 domain (purple) encodes profilin (dark blue). Actin monomers bind *Sc* profilin with an affinity of 2.9 µM (39), and delivery occurs through a loop closure reaction that enables the formin’s FH2 domain (green) to bind the incoming monomer. (B) Architectures of the FH1 domains of Bni1p constructs encoding *Sc* profilin. The FH1 domain is shown to-scale; profilin is not. All polyproline tracts sequences (purple ovals) have been replaced with poly-Gly/Ser sequences matching the length of each tract (white ovals with an X). All constructs also include the FH2 domain, which is located directly C-terminal to the FH1 domain (not shown). (C) Representative inverted fluorescence micrographs depicting the elongation of actin filaments in the presence of Bni1p PRF_38_FH2 or Bni1p PRF_97_FH2. Barbed ends of free (blue) and formin-bound (magenta) filaments are indicated with asterisks. Reactions include 0.75 µM actin monomers (33% Oregon Green-labeled) and 5 nM formin. The scale bar is 5 µm. (D) Barbed end elongation rates mediated by Bni1p FH2 and PRF_xx_FH2 constructs in the presence of 0.75 µM actin monomers (33% Oregon Green-labeled), as seen in panel C. Error bars are standard deviations of the mean elongation rates measured for at least 15 formin-bound filaments visualized in at least 3 independent assays. (E) Representative inverted micrographs collected 300 s after the initiation of polymerization of 0.75 µM actin (33% Oregon Green-labeled) in the presence of 5 nM Bni1p FH2 or PRF_xx_FH2. (F) Number of filaments visualized per 5,000 µm^2^ in micrographs collected 300 s after the initiation of polymerization of 0.75 µM actin monomers in the presence of 5 nM Bni1p FH2 or PRF_xx_FH2, as shown in panel E. Error bars are standard deviations of the mean number of filaments visualized in 3 independent assays.

We assessed the activities of the PRF_xx_FH2 proteins by visualizing actin polymerization in the presence of each variant by TIRF microscopy (Figure 3C). All four PRF_xx_FH2 proteins assemble actin filaments, which are dimmer in fluorescence than filaments with free barbed ends. Profilin binds more weakly to actin monomers labeled at cysteine 374 than to unlabeled monomers (42). Because FH1-mediated polymerization requires profilin-actin complexes, filaments assembled by formins with active FH1 domains undergo biased incorporation of unlabeled subunits (19). The reduction in fluorescence for filaments bound by the PRF_xx_FH2 variants therefore indicates that FH1-tethered profilin competently binds and delivers actin to barbed ends during polymerization.

Consistent with published measurements, filaments bound by the PRF_xx_FH2 proteins elongate at rates ranging from 0.5–1.2 times the rate generated by the FH2 domain (Figure 3D) (34). PRF_22_FH2 mediates the slowest elongation. The elongation rate increases approximately two-fold when FH1-tethered profilin is located 38 residues away from the FH2 domain. The elongation rate continues to increase gradually as the distance separating profilin and the FH2 domain increases.

Fluorescence micrographs collected at the 300-second timepoint contain a larger number of filaments in reactions performed with identical concentrations of each PRF_xx_FH2 construct than with Bni1p FH2 (Figure 3E and F). PRF_38_FH2 assembles the fewest filaments. The nucleation efficiency increases when the FH1-tethered profilin is located either closer to, or farther away from, the FH2 domain. The impact of profilin’s position on nucleation is asymmetrical, as reflected by the much larger number of filaments produced by PRF_22_FH2 than by PRF_97_FH2.

### The FH1 domain transfers profilin-actin to the FH2 dimer during nucleation

The position-dependent contributions of FH1-tethered profilin to filament nucleation and elongation differ significantly (Figure 3D and F), suggesting that FH1-mediated delivery regulates each process differently. To understand how changes in nucleation and elongation collectively impact the net polymerization activity of Bni1p, we monitored the progress of pyrene-actin assembly in the presence of the PRF_xx_FH2 proteins. We found that each variant stimulates polymerization at a rate that exceeds the activity of Bni1p FH2 up to 4-fold (Figure 4A). The rate of polymerization increases continuously as the distance separating profilin and the FH2 domain increases (Figure 4B). The relatively small differences in the elongation activities of Bni1p FH2 and the PRF_xx_FH2 variants suggest that the nucleation properties of these proteins contribute significantly to these differences in bulk polymerization.

**Figure 4.**
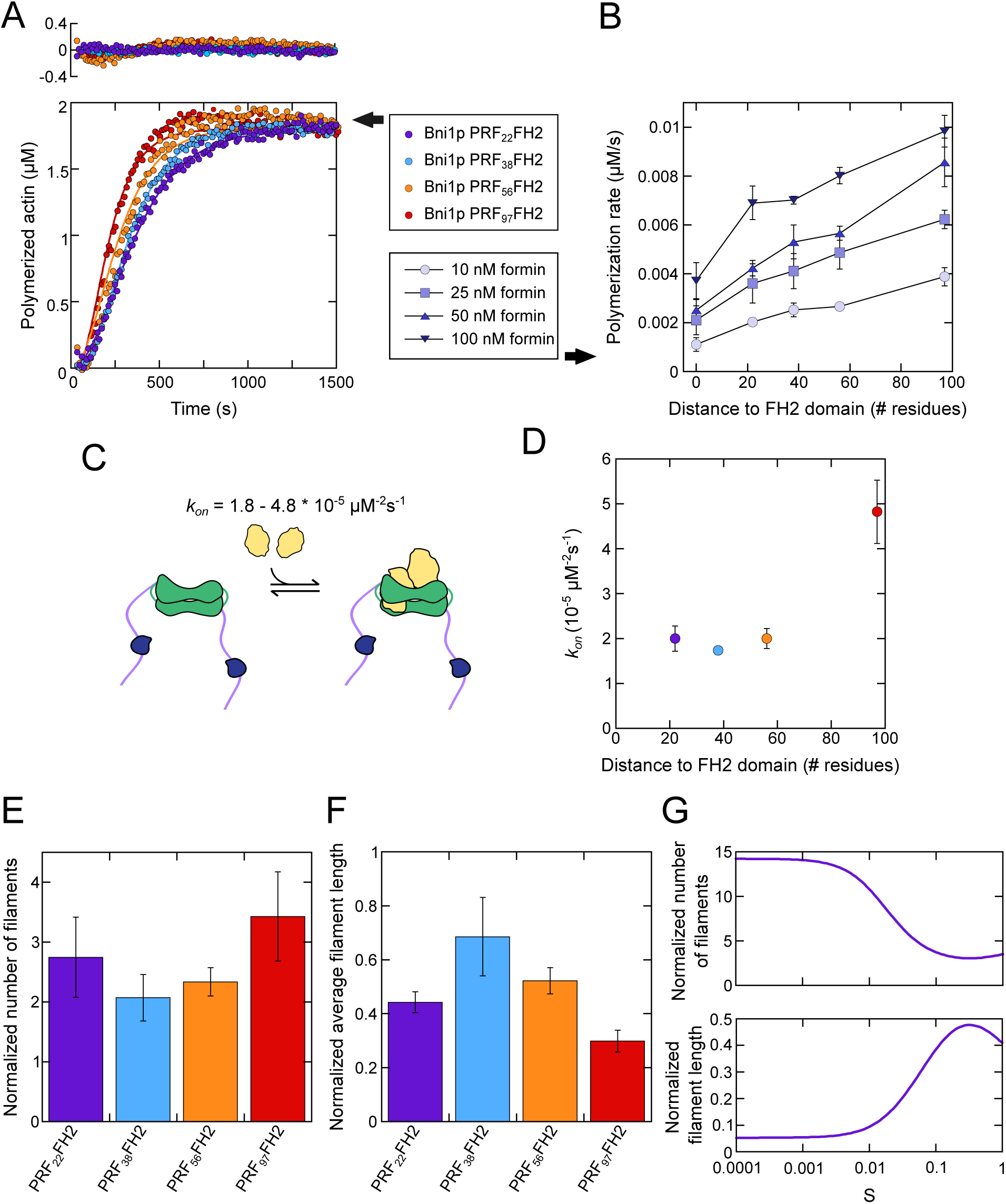
The FH1 domain transfers profilin-actin to the FH2 dimer during nucleation. (A) Representative time courses of spontaneous polymerization of 2 µM actin (20% pyrene-labeled) in the presence of 50 nM Bni1p PRF_xx_FH2 constructs (purple, blue, orange, and red data). For clarity, every fourth data point is plotted. Solid lines are global fits to the data using our kinetic model for PRF_xx_FH2-mediated actin polymerization. Upper panel shows residuals for each fit. (B) Dependence of spontaneous polymerization rates mediated by Bni1p FH2 or Bni1p PRF_xx_FH2 constructs as a function of the distance separating the FH1-tethered profilin from the FH2 domain. Data plotted at a distance of “0 residues” correspond to measurements performed with Bni1p FH2, which does not encode profilin. Polymerization rates were obtained from the slope of the change in fluorescence signal where half of the actin is polymerized in reactions containing 10 nM (circles) 25 nM (squares), 50 (triangles), and 100 nM formin (inverted triangles) formin. Error bars are standard deviations of the mean polymerization rate calculated from at least 3 independent assays. (C) Schematic depicting the assembly of the FH2-actin-actin complex (green and yellow) mediated by a formin whose FH1 domain (purple) encodes profilin (dark blue). The range of association rates extracted from kinetic fits to our experimental data is indicated. (D) Rates of assembly of the FH2-actin-actin complex obtained from kinetic fits to experimental polymerization time courses as a function of the distance separating the FH1-tethered profilin from the FH2 domain. Error bars are standard deviations of the mean association rate extracted from fits to four independent reactions. (E, F) Total number (E) and average equilibrium lengths of filaments (F) assembled by the PRF_xx_FH2 formins obtained from kinetic fits to experimental polymerization time courses as a function of the distance separating the FH1-tethered profilin from the FH2 domain. Both sets of values are normalized to the number and equilibrium lengths of filaments assembled by Bni1p FH2. Error bars are standard deviations of the mean association rates extracted from fits to four independent reactions. (G) Number and equilibrium lengths of filaments assembled by 50 nM PRF_22_FH2 in simulated polymerization reactions as a function of a scaling prefactor (S) that accounts for the diffusional properties of the FH1 domain. A scaling prefactor value of 1 corresponds to an ideal random coil. Both sets of values are normalized to the number and equilibrium lengths of filaments assembled by Bni1p FH2.

We expanded our kinetic model to account for binding of actin monomers to FH1-tethered profilin and FH1-facilitated delivery of actin to FH2-bound nuclei and filament barbed ends (Figures 3A and 4C, Supplemental Table S1). Consistent with previous studies, we assumed that FH1-tethered profilin binds actin monomers and barbed ends with affinities matching those of free profilin, but that the association rate for monomer binding is slower owing to restricted lateral and rotational mobility (34, 35). We also used the number of residues separating FH1-tethered profilin from the FH2 domain to calculate the distance-dependent, intramolecular “loop closure” rate that facilitates FH1-mediated delivery (Figure 3A, Supplemental Table S1) (18).

We used our model to fit our experimental polymerization traces (Figure 4A, solid lines). Each fit produced a trimolecular association rate corresponding to the creation of the FH2-actin-actin species during nucleation. In all reactions containing PRF_xx_FH2 constructs, this association rate exceeds the binding rate that was determined from the polymerization time courses collected with Bni1p FH2 (Figure 4D). This is consistent with a model in which the FH1 domain transfers actin monomers to the FH2 dimer to speed the assembly of filament nuclei. Our model cannot resolve the kinetics of the two individual FH2:actin binding events that lead to the assembly of the FH2-actin-actin species. However, our fits reveal that the FH1-to-FH2 transfer reaction ultimately speeds the formation of this nucleation intermediate 4-to-12-fold, depending on the position of the tethered profilin. PRF_97_FH2 mediates the fastest nucleation, consistent with its efficient bulk polymerization activity (Figure 4A). The nucleation rate displays a biphasic dependence on the distance separating profilin from the FH2 domain, such that the slowest rate is observed at 38 residues of separation.

To determine how FH1-mediated delivery influences the populations of filaments assembled by Bni1p, we used our fitted curves to determine the number of filaments polymerized by each PRF_xx_FH2 variant and their average equilibrium lengths (Figure 4E and F). We found that the number of formin-assembled filaments increases 3.5-fold over the number generated by Bni1p FH2 when profilin is tethered to the FH1 domain 97 residues away from the FH2 domain. This is consistent with the larger numbers of filaments we observed in fluorescence micrographs collected during PRF_97_FH2-mediated polymerization compared to reactions containing Bni1p FH2 (Figure 3E). As the distance separating profilin from the FH2 domain shortens, the number of filaments assembled by the PRF_xx_FH2 variants decreases, until a minimum is reached at 38 residues of separation. At this position, FH1-mediated delivery facilitates the nucleation of twice as many filaments as are polymerized by Bni1p FH2. Filaments assembled by PRF_38_FH2 attain longer average lengths than those assembled when profilin is positioned farther away from the FH2 domain (Figure 4F).

The number of filaments assembled over the course of polymerization increases 50% when the distance separating FH1-tethered profilin from the FH2 domain shortens from 38 to 22. Although consistent with the biphasic dependence on the position of profilin, the magnitude of this effect on nucleation falls short of the nearly 10-fold increase in filament formation observed in our fluorescent micrographs (Figure 3F). This may arise from a restriction in the conformational freedom of the FH1 domain that impacts the rate of delivery when profilin is tethered 22 residues away from the FH2 domain. This would be consistent with a change in the rate of intramolecular loop closure, which has been shown to deviate from random coil predictions for some short polypeptides (43). To test this hypothesis, we varied the “scaling prefactor” which accounts for the diffusional properties of the FH1 domain in our kinetic model. A prefactor value of “1” assumes that the FH1 domain behaves as an ideal random coil (18). In simulated polymerization reactions, we found that the value of this prefactor dramatically influences the number and lengths of the filaments assembled by PRF_22_FH2 (Figure 4G). When the scaling prefactor is set to 0.01, the number of formin-nucleated filaments produced in our simulations is consistent with the relative numbers of filaments observed in our fluorescence micrographs. This is consistent with a model in which a significant reduction in conformational freedom constrains the rate of FH1-mediated delivery and dramatically increases formin-mediated nucleation when profilin binds the FH1 domain at a site near the FH2 domain.

### A gradient of affinities for profilin distributes polymerization activity across the FH1 domain

Our experiments with the PRF_xx_FH2 variants revealed that the location at which actin binds the FH1 domain is a key determinant of the efficiency of formin-directed polymerization. However, whereas FH1-tethered profilin binds actin with equal affinity in each PRF_xx_FH2 construct, the polyproline tracts found in wild-type FH1 domains bind profilin with a wide range of affinities. To determine how variations in profilin binding affinity impact formin activity, we quantified actin polymerization mediated by a series of variants of Bni1p FH1FH2 whose FH1 domains contain a single polyproline tract. We constructed each variant by replacing three of the four polyproline tracts in Bni1p’s FH1 domain with poly-Gly/Ser sequences matching the lengths of the replaced tracts (Figure 5A). As such, each construct retains one polyproline tract, whose sequence and location specify its affinity for profilin and the rate of FH1-mediated delivery (Figure 5B). In a previous study, we named these “single-tract” variants “PY-FH2”, where “PY” refers to the name of the tract that is retained. (30). To facilitate direct comparison with our PRF_xx_FH2 variants, we refer to each single-tract variant as “PY_xx_FH2” here, where “xx” refers to the number of residues separating each polyproline tract from the FH2 domain.

**Figure 5.**
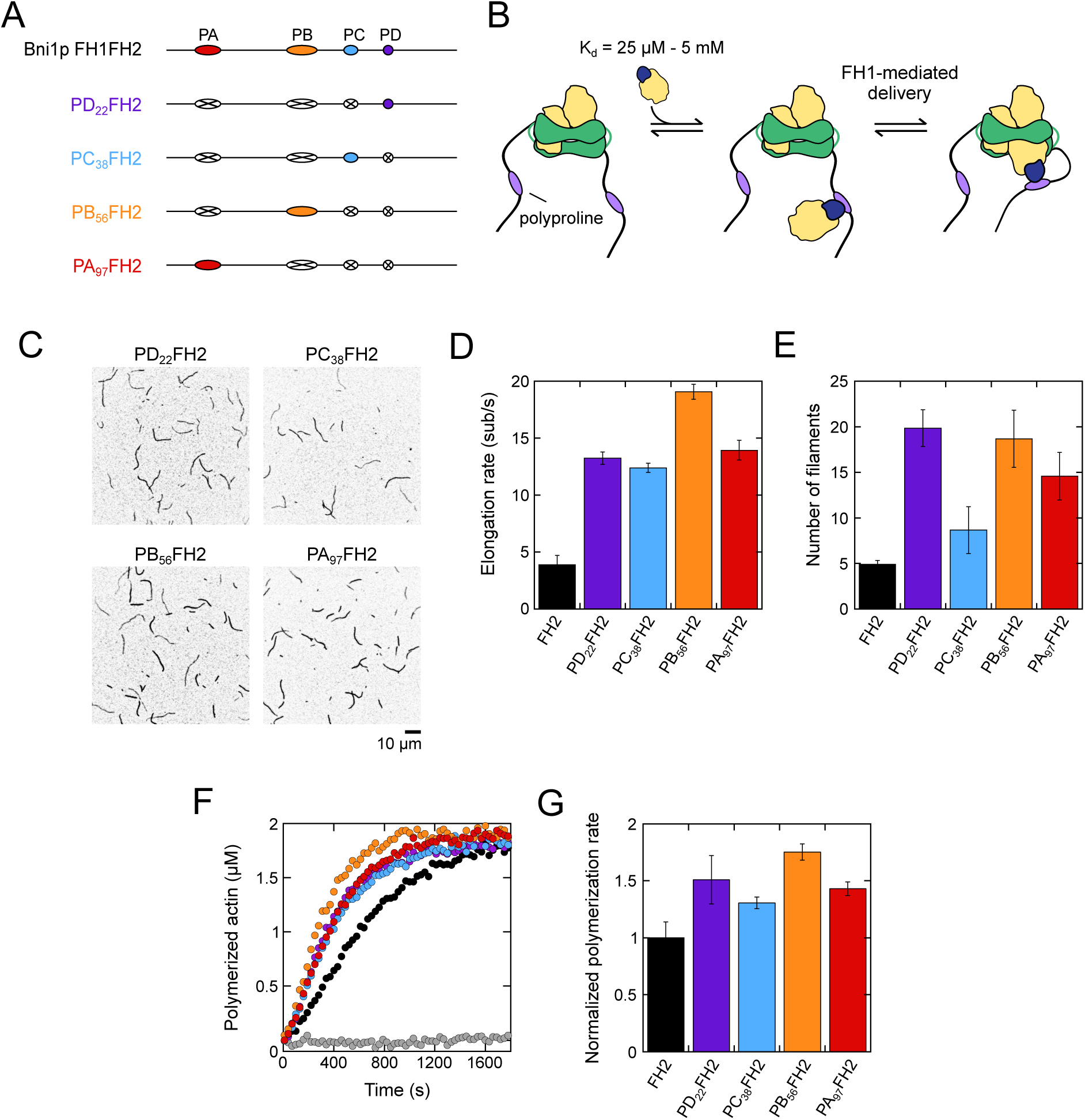
A gradient of affinities for profilin distributes polymerization activity across the FH1 domain. (A) Domain architectures of the FH1 domains of Bni1p constructs encoding single polyproline tracts. The FH1 domain and all polyproline tracts are shown to-scale. In each construct, three polyproline tracts sequences have been replaced with poly-Gly/Ser sequences matching the length of each tract (white ovals with an X). All constructs also include the FH2 domain, which is located directly C-terminal to the FH1 domain (not shown). (B) Schematic depicting the binding and delivery of a profilin-actin complex (blue and yellow) by a formin whose FH1 domain encodes a single polyproline tract (purple). Delivery occurs through a loop closure reaction that enables the formin’s FH2 domain (green) to bind the incoming monomer. (C) Representative inverted micrographs collected 300 s after the initiation of polymerization of 0.75 µM actin (33% Oregon Green-labeled) in the presence of 5 µM profilin and 7.5 nM PY_xx_FH2. (D) Barbed end elongation rates mediated by Bni1p and each PY_xx_FH2 construct in the presence of 0.75 µM actin monomers and 5 µM profilin. Error bars are standard deviations of the mean elongation rates measured for at least 15 formin-bound filaments visualized in at least 3 independent assays. (E) Number of filaments visualized per 5,000 µm^2^ in micrographs collected 300 s after the initiation of polymerization of 0.75 µM actin monomers, 5 µM profilin, and 7.5 nM Bni1p FH2 or PY_xx_FH2 as visualized in panel C. Error bars are standard deviations of the mean number of filaments quantified in 3 independent assays. (F) Representative time courses of spontaneous polymerization of 2 µM actin (20% pyrene-labeled) in the presence of 5 µM profilin and 50 nM Bni1p FH2 (black data) or PY_xx_FH2 constructs (purple, blue, orange, and red data). Gray data were collected in the absence of formin. For clarity, every third data point is plotted. (G) Polymerization rates obtained from the slope of the change in fluorescence signal where half of the actin is polymerized in panel F. Polymerization rates are normalized relative to the activity of the Bni1p FH2 domain. Error bars are standard deviations of the mean polymerization rate calculated from 3 independent assays.

All four single-tract variants promote robust filament assembly in the presence of 5 µM profilin (Figure 5C). Consistent with our previous study, each variant mediates faster filament elongation than does Bni1p FH2, indicating that all four polyproline tracts competently bind and deliver profilin-actin to barbed ends (Figure 5D) (30). Filaments bound by the PB_56_FH2 variant elongate nearly 50% faster than do filaments bound by the other variants. Micrographs collected at the 300-second timepoint show approximately half as many filaments in reactions containing PC_38_FH2 than in reactions containing identical concentrations of the other variants (Figure 5C and E). These differing polymerization activities suggest that the affinity with which a polyproline tract binds profilin differentially modulates the contributions of FH1-mediated delivery to nucleation and elongation.

All four single-tract variants stimulate bulk actin assembly in the presence of 5 µM profilin beyond the rate observed for the Bni1p FH2 domain (Figure 5F and G). In contrast to the PRF_xx_FH2 variants, whose polymerization activities depend on the position of the FH1-tethered profilin, the polymerization rates mediated by the single-tract variants vary in a position-independent manner (Figure 5C). Therefore, variations in profilin-binding affinity across the ensemble of polyproline tracts contained in the FH1 domain facilitate engagement of each tract during polymerization. As revealed by our TIRF micrographs, the precise contributions made by each polyproline tract to filament nucleation and elongation differ depending on the sequence and position of the tract.

## Discussion

Formins promote the assembly of unbranched actin filaments through direct and indirect interactions with actin monomers and filament ends (10, 14-16). To understand how the actin-binding protein profilin influences formin activity, we analyzed the mechanism of filament nucleation mediated by the *S. cerevisiae* formin Bni1p. We found that the FH1 domain speeds *de novo* filament assembly by transferring actin to the FH2 domain at a rate that depends on the locations of its polyproline tracts and the affinity of each tract for profilin.

### FH2 dimers bind profilin-actin inefficiently

Previous studies proposed that FH2-mediated nucleation proceeds via the initial formation of an FH2-actin-actin complex (20, 33). Because binding of the second actin monomer to the complex is rate-limiting, it was unknown whether FH2 binding requires actin dimerization, or if actin monomers can bind to FH2 dimers sequentially. By applying a deterministic model to fit experimental time courses of polymerization, we quantified the rate of formation of the FH2-actin-actin species (Figure 1A). Two sets of widely divergent kinetic parameters defining spontaneous actin dimerization and trimerization produced identical rates of formin-mediated nucleation, indicating that the formation of the FH2-actin-actin complex does not require actin dimerization. Rather, FH2 domains can bind individual monomers sequentially. These results reveal that formin-mediated and spontaneous filament nucleation are parallel processes that consume the same pool of actin but do not otherwise intersect.

In the presence of profilin, preferential binding of actin monomers over profilin-actin complexes slows filament nucleation (Figure 2C). In contrast, FH2-bound filament ends incorporate actin monomers and profilin-actin complexes with equal efficiency at the concentrations of profilin we tested (29). The preference for actin monomers during nucleation suggests that the nature of the interactions between FH2 domains and actin differs during the nucleation and elongation processes. Recent cryo-EM structures of formin-bound filaments provide some insights into these differences (15). These structures revealed that FH2 domains and profilin bind overlapping surfaces on the terminal actin subunit at the barbed end. This structural overlap may sterically hinder the binding of profilin-actin complexes to free FH2 dimers during nucleation. Following the assembly of a filament nucleus, the newly-created barbed end might then serve as a template that orients incoming profilin-actin complexes in a way that is compatible with FH2 domain binding during elongation. Rapid dissociation of profilin from newly-polymerized actin subunits ultimately resolves steric clashes with the FH2 domain, enabling continued subunit addition (25).

### FH1-mediated actin delivery differentially regulates nucleation and elongation

By comparing the activities of Bni1p FH2 and Bni1p FH1FH2, we found that the FH1 domain speeds polymerization in the presence of profilin by stimulating both nucleation and elongation (Figure 2). The impact on nucleation is consistent with a model in which FH1-mediated delivery facilitates the binding of profilin-actin complexes to the FH2 domain (Figure 6). The process of delivery occurs through an intramolecular “loop closure” reaction, which brings two sections of a flexible polypeptide chain into contact (18). In this manner, FH1-mediated delivery increases the frequency of collisions between profilin-actin complexes and FH2 dimers, thereby promoting binding. Consistent with the disordered nature of FH1 domains, the loop closure reaction is diffusion-limited (18). As a result, the rate of delivery slows as the distance between the profilin binding site and the FH2 domain increases (27).

**Figure 6.**
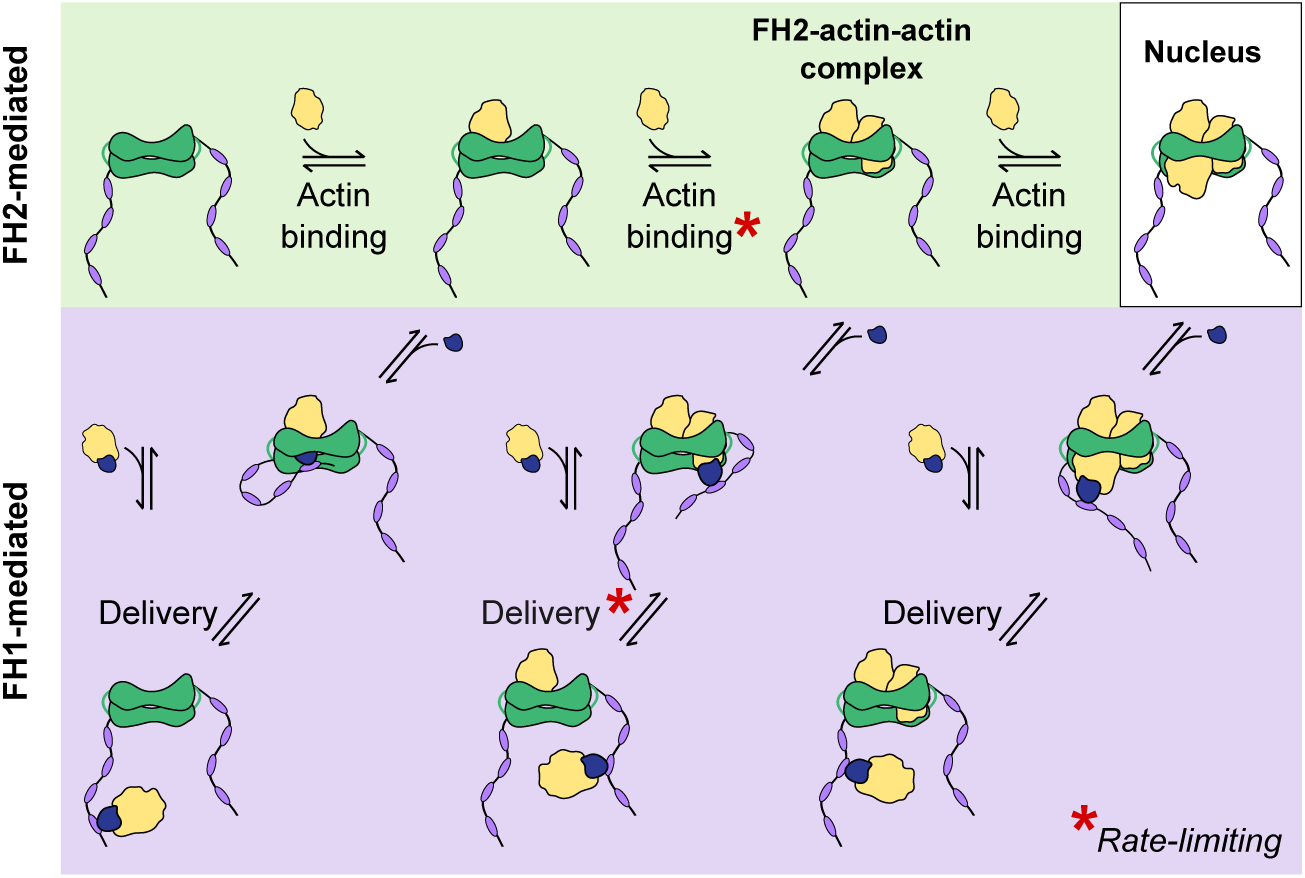
Model for formin-mediated filament nucleation. Schematic representation of actin filament nucleation mediated by the FH1 (purple) and FH2 (green) domains of Bni1p. Top, The FH2 dimer binds actin monomers (yellow) in three sequential events. The second monomer binding event is rate-limiting and results in the formation of the FH2-actin-actin complex. Binding a third monomer results in a formin-bound filament nucleus. Bottom, Profilin-actin complexes (yellow and blue) bind polyproline tracts (purple) in the FH1 domain, facilitating their delivery to the FH2 dimer. Transfer of the second monomer to the FH2 dimer is rate-limiting.

By tethering profilin at different positions in the FH1 domain, we probed the effects of varying the loop closure rate on the mechanism of formin-mediated polymerization. We found that the rate of polymerization increases as the distance separating the FH1-tethered profilin and the FH2 domain increases, revealing an inverse-dependence on the loop closure rate. Notably, this effect on the overall polymerization activity arises from position-specific modulation of the efficiency of filament nucleation and elongation (Figure 3D and F). The precise impacts on nucleation and elongation differ, suggesting that FH1-mediated delivery contributes differently to each process.

In addition to promoting actin binding and delivery, covalent attachment of profilin to the FH1 domain introduces a local concentration of profilin relative to the FH2 domain. This local concentration depends on the position at which profilin is tethered and determines the probability of profilin binding to the terminal actin subunit at an FH2-bound barbed end. Because profilin binding sterically inhibits actin monomer addition, filament elongation slows as the local concentration of profilin increases (34). Consistent with a decrease in the local concentration of profilin, the elongation rate mediated by the PRF_xx_FH2 variants increases as the number of residues separating profilin and the FH2 domain increases (Figure 3D). Therefore, occupancy of the barbed end by profilin limits the contributions of FH1-mediated delivery to elongation.

In contrast to the monophasic effect on elongation, variation of the loop closure rate produces a biphasic effect on formin-mediated nucleation. Filament nucleation is least efficient when profilin is tethered 38 residues away from the FH2 domain. Increasing this distance to 56 and 97 residues boosts the number of formin-assembled filaments 3- and 5-fold, consistent with distance-dependent changes in the local concentration of profilin. The nucleation activities of these PRF_xx_FH2 variants span a much larger range than do their elongation activities, which differ by less than 20%. Therefore, the probability with which profilin binds to barbed ends regulates the assembly of filament nuclei to a greater extent than it affects subunit addition during filament elongation. Given their overlapping binding sites on the actin surface, profilin binding may induce the dissociation of actin monomers from FH2 dimers, thereby driving the disassembly of nascent filament nuclei. Such a reversal of nuclear assembly would magnify the effects of profilin binding on the rate of *de novo* filament formation. The large range of nucleation activities displayed by the PRF_xx_FH2 variants also suggests that FH1-mediated delivery stimulates the assembly of FH2-actin-actin complexes to an extent that exceeds its contributions to filament elongation. This is consistent with a role for the FH1 domain in positioning incoming profilin-actin complexes in a way that overcomes steric clashes and facilitates their binding to the FH2 domain during nucleation.

Filament nucleation is most efficient when profilin is tethered 22 residues away from the FH2 domain, indicating that rapid loop closure can compensate for a high local concentration of profilin to drive the assembly of FH2-actin-actin complexes. A kinetic model that assumes the FH1 domain behaves as a random coil fails to capture the magnitude of nucleation mediated by PRF_22_FH2 (Figures 3F and 4E). A reduction in the scaling prefactor accounting for the diffusional properties of the FH1 domain dramatically increases the nucleation efficiency in simulated polymerization reactions (Figure 4G). Therefore, it is likely that profilin uniquely restricts the conformational freedom of the FH1 domain when bound at a site near the FH2 domain. Such a diffusional constraint might speed the transfer of profilin-actin complexes to empty FH2 dimers. Alternatively, a reduction in conformational sampling might restrict the position of FH1-tethered profilin in a way that prevents its binding to the FH2-actin-actin complex, thereby increasing the stability of nascent filament nuclei.

### The FH1 domain differentially engages its polyproline tracts to maximize the efficiency of polymerization

In contrast to our PRF_xx_FH2 variants, which bind actin with uniform affinity, the polyproline tracts encoded in wild-type FH1 domains bind profilin-actin complexes with micromolar-to-millimolar affinities. The FH1 domains of many formins are organized in a way that positions the shortest polyproline tracts (which bind profilin weakly) closest to the FH2 domain and the longest tracts (which bind profilin more tightly) farther away (27). This establishes a gradient of affinities for profilin that is coordinated with distant-dependent rates of FH1-mediated delivery to maximize the efficiency of filament elongation (27).

Using a series of constructs of Bni1p containing single polyproline tracts at their native locations in the FH1 domain, we determined that variations in profilin-binding affinity along the FH1 domain minimize the differences in the rate of polymerization mediated by each polyproline tract. Whereas the net polymerization activities of the PRF_xx_FH2 variants vary up to 200% depending on the position at which profilin is tethered to the FH2 domain, the rate of actin assembly mediated by the single-tract constructs varies by only ∼35%. Notably, individual tracts possess specialized nucleation and elongation activities. It is therefore likely that certain tracts contribute primarily to *de novo* filament assembly, whereas others play a more prominent role during elongation. Collectively, the unique set of polyproline tract locations and sequences found within FH1 domains maximize the efficiency of formin-mediated polymerization through optimized engagement during polymerization.

### Specialized mechanisms regulate formin-mediated polymerization in cells

By dissecting the contributions of the FH1 and FH2 domains to formin-mediated polymerization, we have revealed a general mechanism for actin assembly in the presence of profilin. As all formins possess an FH1 and an FH2 domain, this mechanism likely captures a set of interactions that are universal to all formins (44, 45). Beyond the basal functions of these two domains, additional cellular factors further tune the nucleation and elongation activities of formins. Importantly, the C-terminal tail domains of some formins have been shown to contribute to filament assembly through interactions with actin monomers or along the lengths of filaments (46). In *S. cerevisiae*, the myosin passenger protein Smy1p binds the formin Bnr1p and inhibits its actin polymerization activity (47). Binding of Spire, Ena/Vasp, and adenomatous polyposis coli regulates the nucleation activities of other formins (48-50). A comprehensive picture of the cellular activities of formins will therefore require mapping of these complex and dynamic interactions networks.

## Materials and Methods

### Protein purification

Constructs encoding the FH1 and FH2 domains of Bni1p (“Bni1p FH2”: residues 1329-1766; “Bni1p FH1FH2”: residues 1228-1766) and variants thereof were cloned into pGEX-4T-3 plasmids (GE Healthcare). Each construct encoded a TEV protease recognition sequence following the N-terminal GST tag and a C-terminal 6-His tag. A series of “single-tract” variants was designed by substituting all but one of the polyproline tracts in the FH1 domain of the “Bni1p FH1FH2” construct with a repeated Gly-Gly-Gly-Ser sequence matching the length of each replaced tract. A series of chimeric profilin-formin constructs was generated by replacing the remaining polyproline tract in each single-tract variant with the nucleotide sequence encoding *S. cerevisiae*. An additional linker sequence encoding the amino acids Gly-Gly-Ser was included following the profilin sequence to promote flexibility. To enhance protein stability, each PRF_xx_FH2 variant was N-terminally truncated to include only 10-28 residues preceding the profilin sequence. These hybrid profilin-formin constructs were named “PRF_xx_FH2”, where “xx” corresponds to the number of residues in the FH1 domain separating profilin from the FH2 domain. The Bni1p residue boundaries for the PRF_xx_FH2 variants are as follows: 1228-1766 (Bni1p PRF_97_FH2), 1251-1766 (Bni1p PRF_56_FH2), 1292-1766 (Bni1p PRF_38_FH2), 1310-1766 (Bni1p PRF_22_FH2).

All Bni1p variants were expressed overnight at 16 °C in 1 L cultures of BL21 DE3 RP Codon Plus cells (Agilent Technologies). Cell pellets were resuspended in 50 mM Tris (pH 8.0), 500 mM NaCl, 1 mM DTT, and lysed by sonication. Clarified lysates were incubated with 2 mL glutathione-sepharose resin (Gold Biotechnology) with rotation for 1 hour at 4 °C. Resin and protein solutions were transferred to empty chromatography columns, washed with 10 volumes of lysis buffer, followed by 10 volumes of low-salt wash buffer (50 mM Tris (pH 8.0), 100 mM NaCl, 1 mM DTT), and eluted with 3 volumes of 100 mM glutathione (pH 8.0) in low-salt wash buffer. Glycerol was added to Bni1p PRF_xx_FH2 eluates to a final concentration of 10% (v/v). Each protein was incubated with 5 µM TEV protease overnight at 4 °C to remove the GST tag. Formins were separated from cleaved GST and the TEV protease by nickel affinity chromatography, concentrated using 30,000 MWCO spin columns (Millipore Sigma) and dialyzed into KMEI (50 mM KCl, 1 mM MgCl2, 1 mM EGTA, 10 mM imidazole (pH 7.0)), with 1 mM DTT. Glycerol was included in the dialysis buffer at a concentration of 10% (v/v) for PRF_xx_FH2 proteins. Purified proteins were flash-frozen in liquid nitrogen and stored at -80 °C. We used ProtParam (www.expasy.org/protparam) (51) to calculate extinction coefficients for all constructs.

S. cerevisiae profilin was expressed from a pMW172 vector in BL21 DE3 pLysS cells and purified as previously described (29). We used an extinction coefficient of 19,060 M^-1^cm^-1^ at λ = 280 nm to determine the concentration of profilin. Skeletal muscle actin was purified from an acetone powder prepared from frozen chicken breasts (Trader Joe’s) via one cycle of polymerization and depolymerization (52). Actin monomers were polymerized by dialyzing in 100 mM KCl, 2 mM MgCl2, 25 mM Tris-HCl (pH 7.5), and 0.3 mM ATP, and incubated at 4 °C overnight with a 1:10 molar ratio of actin to pyrenyl iodoacetamide (Thermo Fisher Scientific) or Oregon Green maleimide (Thermo Fisher Scientific). Labeled and unlabeled actin monomers were gel-filtered on S-300 resin (GE Healthcare) in G-Buffer (2 mM Tris (pH 8.0), 0.2 mM ATP, 0.5 mM DTT, 0.1 mM CaCl2) and stored at 4 °C. We used extinction coefficients of 26,000 M^-1^cm^-1^ at λ = 290 nm for unlabeled actin, 22,000 M^-1^cm^-1^ at λ = 344 nm for pyrene, and the following relation to calculate the concentration of pyrene-labeled actin: [total actin] = (A_290_ – (A_344_*0.127))/26,000 M^-1^cm^-1^. We used an extinction coefficient of 78,000 M^-1^cm^-1^ at λ = 491 for Oregon Green, and the following relation to calculate the concentration of Oregon Green-labeled actin: [total actin] = (A_290_ – (A_491_*0.171))/26,000 M^-1^cm^-1^.

### Pyrene-actin assembly assays

Time courses of actin polymerization were collected by measuring pyrene fluorescence emission with a Molecular Devices SpectraMax Gemini EM fluorescence plate reader using Corning 96-well flat-bottom plates. Reactions containing 2 µM actin monomers (20% pyrene-labeled) were polymerized in the absence or presence of a range of concentrations of formin and profilin in 10 mM imidazole (pH 7.0), 50 mM KCl, 1 mM MgCl_2_, 1 mM EGTA, 0.17 mM ATP, 0.5 mM DTT, 0.03 mM CaCl2, and 0.17 mM Tris-HCl (pH 8.0). Samples were excited at 365 nm and the fluorescence emission intensity was measured every 10 s at 407 nm over a period of 30-60 min. The fluorescence signal was converted to polymer concentration by normalizing the fluorescence intensity to the final predicted actin polymer concentration, assuming a critical concentration of 0.17 µM in the absence of profilin (7). Bulk polymerization rates were calculated from the slopes of linear fits applied to the polymerization time courses at the point where half of the actin is polymerized.

### Microscopy and data analysis

Glass slides and coverslips (22 x 50 mm; Thermo Fisher Scientific) were washed and assembled into open-ended flow chambers as described previously (29). Chambers were prepared via sequential incubations with 0.5% Tween 20 in High-Salt TBS (HS-TBS; 50 mM Tris (pH 7.5), 600 mM NaCl), ∼ 1 µM N-ethylmaleimide-inactivated chicken skeletal muscle myosin (53) in HS-TBS, and 100 mg/mL bovine serum albumin (BSA) in HS-TBS. Chambers were washed with HS-TBS following each incubation step and with KMEI prior to the introduction of a polymerization reaction.

Solutions of unlabeled and labeled Ca^2+^-ATP actin monomers were converted to Ca2+-ATP actin monomers via the addition of 50 µM MgCl2 and 0.2 mM EGTA. Polymerization was initiated by the addition of 2x microscopy buffer (1x microscopy buffer: 10 mM imidazole (pH 7.0), 50 mM KCl, 1 mM MgCl_2_, 1 mM EGTA, 50 mM DTT, 0.2 mM ATP, 15 mM glucose, 20 µg/mL catalase, 100 µg/mL glucose oxidase, 0.5% (w/v) methylcellulose (4,000 cP at 2%)) in the absence or presence of formin and profilin. Reactions were introduced into a flow chamber for imaging immediately upon initiation of polymerization.

Polymerization reactions were visualized by through-objective TIRF microscopy on an Olympus Ti83 motorized microscope outfitted with a CellTIRF system using a 60x 1.49 N.A. objective and a 488-nm laser. Time-lapse images were collected every 10 s using a Hamamatsu C9100-23B ImagEM X2 EMCCD camera and CellSens Dimension software (Olympus; https://www.olympus-lifescience.com/en/software/cellsens/). Images were processed with ImageJ software (National Institutes of Health; https://imagej.net/software/fiji/) (54). For elongation measurements, changes in length for at least 15 formin-bound filaments and 10 control (i.e., not formin-bound) filaments were measured over a span of at least 200 s. Linear fits were applied to plots of filament length over time using Kaleidagraph software (Synergy Software; https://www.synergy.com/). For nucleation measurements, the number of filaments present in 5,000 µm^2^ or 10,000 µm^2^ fields were counted.

### Kinetic modeling

Kinetic schemes described by Rosenbloom et al, Sept et al, Vavylonis et al, Paul and Pollard, and Zweifel and Courtemanche (5, 18, 20, 34, 37) were combined to generate a single model of actin polymerization that includes the nucleation and elongation activities of formins in the absence and presence of profilin. Consistent with published studies (5, 18, 20, 35), reaction parameters for ATP hydrolysis are not explicitly defined in our model but can be accounted for via adjustment of elongation rate constants (18). Experimental polymerization time courses were fitted using the Levenberg-Marquardt algorithm (55, 56) implemented in the Parameter Estimation module of the program COPASI (57) using published rate constants (18, 20, 35). Simulated time courses were calculated using the Time Course module of COPASI in deterministic mode using the LSODA algorithm (58, 59) at fixed bulk concentrations of actin and formin.

## Supporting Information

This article contains supporting information.

