## Supplemental Table 1 for "Mechanism of formin-mediated filament nucleation from profilin-actin"

| **Supplemental Table S1. Kinetic model parameters** | | | | |
|---|---|---|---|---|
| **Reaction** | **Reaction Description** | | **Forward rate** | **Reverse rate** |
| Profilin-Actin Equilibrium | | | | |
| 1 | Profilin binds actin[a] | P + A = PA | 30 $\mu M^{-1}s^{-1}$ | 90 $s^{-1}$ |
| Spontaneous Actin Polymerization | | | | |
| *Spontaneous actin nucleation* | | | | |
| 2 | Monomer dimerization[b] | A + A = 2A | 35.7 $\mu M^{-1}s^{-1}$<br>2.9*$10^6$ $\mu M^{-1}s^{-1}$ | 1.63*$10^8$ $s^{-1}$<br>0.054 $s^{-1}$ |
| 3 | Monomer trimerization[b] | 2A + A = 3A | 2.18 $\mu M^{-1}s^{-1}$<br>2.3*$10^{-4}$ $\mu M^{-1}s^{-1}$ | 1300 $s^{-1}$<br>12 $s^{-1}$ |
| 4 | Free filament formation (addition of A)[b] | 3A + A = Free_BE | 11.6 $\mu M^{-1}s^{-1}$ | 1.4 $s^{-1}$ |
| 5 | Free filament formation (addition of PA)[c] | 3A + PA = Free_BE-P | 10 $\mu M^{-1}s^{-1}$ | Detailed balance |
| *Spontaneous filament elongation* | | | | |
| 6 | Actin incorporation at free barbed end[d] | Free_BE + A = Free_BE + FA | 11.6 $\mu M^{-1}s^{-1}$ | 1.4 $s^{-1}$ |
| 7 | Profilin-actin incorporation at free barbed end[e] | Free_BE + PA = Free_BE-P + FA | 10 $\mu M^{-1}s^{-1}$ | Detailed balance |
| 8 | Actin incorporation at free pointed end[d] | Free_PE + A = Free_PE + FA | 1.3 $\mu M^{-1}s^{-1}$ | 0.8 $s^{-1}$ |
| *Profilin binds to free barbed ends* | | | | |
| 9 | Profilin binds to free barbed end[f] | Free_BE + P = Free_BE-P | 10 $\mu M^{-1}s^{-1}$ | 2500 $s^{-1}$ |
| Formin-Mediated Polymerization | | | | |
| *Formin-mediated nucleation* | | | | |
| 10 | Formin-mediated nucleation[g] *(Assembly of FH2-actin-actin complex)* | FH2 + A + A = Bound_BE | Fitted parameter | 0 |
| 11 | Formin-mediated nucleation (A + PA)[g] *(Assembly of FH2-actin-actin complex from 1 actin monomer and 1 profilin-actin complex)* | FH2 + A + PA = Bound_BE-P | Fitted parameter | 0 |
| 12 | Formin-mediated nucleation (PA + PA)[g] | FH2 + A + PA = Bound_BE-P + P | Fitted parameter | 0 |

| | | | | |
|---|---|---|---|---|
| | *(Assembly of FH2-actin-actin complex from 2 profilin-actin complexes)* | | | |
| 13 | Formin-mediated nucleation[g] *(FH1-assisted assembly of FH2-actin-actin complex)* | $PRF_{xx}FH2$ + A + A = Bound_BE | Fitted parameter | 0 |
| *Profilin binds to formin-bound barbed ends* | | | | |
| 14 | Profilin binds to bound barbed end[f] | Bound_BE + P = Bound_BE-P | 10 $\mu M^{-1}s^{-1}$ | 2500 $s^{-1}$ |
| *FH2-mediated filament elongation* | | | | |
| 15 | Actin incorporation at bound barbed end[h] | Bound_BE + A = Bound_BE + FA | p*11.6 $\mu M^{-1}s^{-1}$ | p*1.4 $s^{-1}$ |
| 16 | Profilin-actin incorporation at bound barbed end[i] | Bound_BE + PA = Bound_BE-P + FA | p*10 $\mu M^{-1}s^{-1}$ | Detailed balance |
| *FH1-mediated filament elongation* | | | | |
| 17 | Actin binds to FH1-tethered profilin[j] | FH1-P + A = FH1-PA | 20 $\mu M^{-1}s^{-1}$ | 60 $s^{-1}$ |
| 18 | FH1-profilin loop closure[k] | FH1-P_o = FH1-P_c | $(S*n_r)^{-3/2}$ $s^{-1}$ | 2500 $s^{-1}$ |
| 19 | FH1-profilin-actin loop closure[l] | FH1-PA_o = FH1-P_c + FA | $p*E_0*(S*n_r)^{-3/2}$ $s^{-1}$ | Detailed balance |
| 20 | Free actin incorporation at pointed end of formin-bound filament[d] | Bound_PE + A = Bound_PE + FA | 1.3 $\mu M^{-1}s^{-1}$ | 0.8 $s^{-1}$ |
| *Pointed end elongation of formin-bound filaments* | | | | |
| 21 | Free actin incorporation at pointed end of formin-bound filament[d] | Bound_PE + A = Bound_PE + FA | 1.3 $\mu M^{-1}s^{-1}$ | 0.8 $s^{-1}$ |

| General Reaction Parameters | | |
|---|---|---|
| Profilin-actin dissociation constant[m] | $K_d$_PA | 3 μM |
| Polymerization prefactor[n] | p | 0.5 (0.25 for $PRF_{22}FH2$) |
| Energetic prefactor[o] | $E_0$ | $10^5$ |
| Scaling prefactor[p] | S | 1 (Variable in simulations of $PRF_{22}FH2$-mediated polymerization) |
| Distance (in residues) from FH1-tethered profilin to FH2 domain | $n_r$ | 22-97 |

[a]The association rate was used in simulations performed by Paul and Pollard (1) and estimated from Vinson and coworkers (2). The dissociation rate was calculated based on the dissociation constant of 3 μM for actin and *S. cerevisiae* profilin (3).

[b]The two sets of widely divergent rates were reported by Sept and McCammon (4) and Rosenbloom et al (5). A filament is established via the association of an actin monomer with an actin trimer, and we treat this reaction as irreversible, consistent with Sept and McCammon and Paul and Pollard (1, 6).

[c]The association rate was used in the simulations performed by Vavylonis and coworkers and Paul and Pollard (1, 7). We treat this reaction as irreversible, consistent with the studies of Sept and McCammon and Paul and Pollard (1, 6).

[d]Rates measured by Pollard (8).

[e]These rates were used by Vavylonis and colleagues (7). The dissociation rate was calculated based on detailed balance.

[f]These rates were estimated by Vavylonis and colleagues and Paul and Pollard (1, 7). The dissociation rate was calculated based on the affinity of human profilin 1 for the barbed ends of filaments (~250 μM (9, 10)) and assumes that the FH2 domain does not influence this affinity (7).

[g]As in the Paul study and our previous work, we assume that formin irreversibly nucleates a filament by assembling an FH2-actin-actin complex. This can occur through binding two actin monomers, one actin monomer and one profilin-actin complex, or two profilin-actin complexes (1, 11). Although formin likely binds the actin monomers and/or profilin-actin sequentially, binding the second actin is rate-limiting. Therefore, we used our kinetic model to fit the association rate for the trimeric FH2-actin-actin complex. When fitting polymerization reactions containing either profilin or the $PRF_{xx}FH2$ constructs, we fixed the rate at which the FH2 binds two actin monomers to the rate obtained from global fits to reactions containing only actin monomers and Bni1p FH2 (i.e., $3.9*10^{-6}$ $\mu M^{-2}s^{-1}$; see Figure 1).

[h]Association and dissociation rates derived from Pollard (8) and multiplied by the polymerization prefactor "p", which reflects the degree to which the FH2 domain slows binding of actin monomers to the barbed end (7).

[i]These rates were used by Vavylonis and colleagues.

[j]The association rate was estimated by Vavylonis and colleagues (7). It was selected based on published rates of 35-53 $\mu M^{-1}s^{-1}$ for the associations of amoeba profilin II and bovine spleen profilin with actin (2, 12), and accounts for the reduced mobility of FH1-bound profilin. Our calculation of the dissociation rate assumes that covalently tethering profilin to the FH1 domain does not influence its affinity for actin.

[k]This rate was used by Vavylonis and colleagues. It assumes the FH1 domain is a random coil and that the FH2 domain does not impede binding of FH1-bound profilin to the barbed end (7).

[l]This rate was used by Vavylonis and colleagues and accounts for the effects of the FH2 domain on the availability of formin-bound barbed ends for actin monomer binding (7).

[m]This dissociation constant was measured by Eads and coworkers (3).

[n] This value was first reported by Kovar and colleagues, who showed that filaments bound by Bni1p FH2 elongate at half the rate observed for filaments with free barbed ends (13). We use the same value for $PRF_{38}FH2$, $PRF_{56}FH2$, and $PRF_{97}FH2$, which mediate approximately the same elongation rate as Bni1p FH2. Filaments bound by $PRF_{22}FH2$ elongation approximately half as fast as those bound by Bni1p FH2, so we use a value of 0.25 when fitting polymerization reactions containing this protein.

[o]This energetic prefactor is a microscopic rate into which all orientational effects have been collapsed (7). The rate of association of the two ends of an unfolded peptide is $10^7$ $s^{-1}$ (14). Our rate is slower to account for the probability of collisions between FH1-tethered profilin and the barbed end occurring in the correct orientation to promote a binding event.

[p]This scaling parameter accounts for FH1 diffusional properties that deviate from those of an ideal random coil.
